# Structural and functional MRI data differentially predict chronological age and behavioral memory performance

**DOI:** 10.1101/2022.03.24.485603

**Authors:** Joram Soch, Anni Richter, Jasmin M. Kizilirmak, Hartmut Schütze, Hannah Feldhoff, Larissa Fischer, Lea Knopf, Matthias Raschick, Annika Schult, Emrah Düzel, Björn H. Schott

**Author notes:** These authors contributed equally to this work (Joram Soch, Anni Richter). **Address for correspondence:** Dr. Joram Soch, German Center for Neurodegenerative Diseases, Von-Siebold-Str. 3a, 37075 Göttingen, Germany, /, PD Dr. Dr. Björn Hendrik Schott, Leibniz Institute for Neurobiology, Brenneckestr. 6, 39118 Magdeburg, Germany, /.

## Abstract

Human cognitive abilities decline with increasing chronological age, with decreased explicit memory performance being most strongly affected. However, some older adults show “successful aging”, that is, relatively preserved cognitive ability in old age. One explanation for this could be higher brain structural integrity in these individuals. Alternatively, the brain might recruit existing resources more efficiently or employ compensatory cognitive strategies. Here, we approached this question by testing multiple candidate variables from structural and functional neuroimaging for their ability to predict chronological age and memory performance, respectively. Prediction was performed using support vector machine (SVM) classification and regression across and within two samples of young (N = 106) and older (N = 153) adults. The candidate variables were (i) behavioral response frequencies in an episodic memory test, (ii) recently described fMRI scores reflecting preservation of functional memory networks, (iii) whole-brain fMRI contrasts for novelty processing and subsequent memory, (iv) resting-state fMRI maps quantifying voxel-wise signal fluctuation and (v) gray matter volume estimated from structural MR images. While age group could be reliably decoded from all variables, chronological age within young and older subjects was best predicted from gray matter volume. In contrast, memory performance was best predicted from task-based fMRI contrasts and particularly single-value fMRI scores, whereas gray matter volume has no predictive power with respect to memory performance in healthy adults. Our results suggest that superior memory performance in healthy older adults is better explained by efficient recruitment of memory networks rather than by preserved brain structure.

## 1. Introduction

Episodic memory performance peaks in young adulthood and declines with increasing age. Notably, a subpopulation of older adults show “successful aging”, with memory performance comparable to that of younger adults (Nyberg et al., 2012; Nyberg and Pudas, 2019). An early assessment of changes in cognitive performance can help to determine people at risk of pathological aging, such as various forms of dementia, and allows for early medical and behavioral interventions (Naismith et al., 2009; Cabeza et al., 2018; Whitty et al., 2020). Machine learning-based techniques such as support vector machine (SVM) classification and regression provide promising approaches to differentiate normal from pathological neurocognitive aging. They have been employed to predict chronological age from structural magnetic resonance imaging (MRI; Cole et al., 2017, 2018), to estimate brain age (Bashyam et al., 2020; Habes et al., 2021) or to distinguish health from disease (Dyrba et al., 2021; Eitel et al., 2021).

In contrast to the abundant literature on age prediction from structural MRI (Cole et al., 2017, 2018; Luders et al., 2016; Steffener et al., 2016; Soch, 2020), few studies have been devoted to predicting cognitive function, particularly memory performance, from neuroimaging data. One such study found that a combination of ApoE genotype and functional MRI was the most effective predictor for future cognitive decline (Woodard et al., 2010). The wide range of cognitive functioning even within narrowly defined age groups suggests that chronological age and cognitive performance might be predicted by different modalities. Several studies evaluated potential structural, functional, physiological and behavioral predictors of age-related cognitive decline (Gross et al., 2011; Hou et al., 2020; Chen et al., 2021), but only few studies systematically compared different predictors and their joint predictive value (e.g., Woodard et al., 2010).

Comparing the predictive value of MRI biomarkers for chronological age versus individual memory performance appears to be a promising endeavor, because “successful aging” may reflect dissociable neural mechanisms: differences in the manifestation of age-related physiological changes (“brain maintenance”) and/or differences in cognitive processing (“cognitive reserve”; Nyberg et al., 2012). Thus, data from different modalities may differentially predict chronological age and memory performance, respectively.

We compared SVM-based prediction of chronological age versus prediction of memory performance from behavioral data, task-based fMRI, resting-state fMRI, and structural MRI markers associated with increasing age. Our analyses where based on a large sample of 106 young and 153 older subjects (Soch et al., 2021a). Episodic memory performance was measured in the fMRI task and in various neuropsychological tests, using either incidental or intentional memory formation.

In addition to task-based fMRI, we also included recently described single-value fMRI scores (Soch et al., 2021b; Richter et al., 2022). These scores are derived from fMRI contrasts and describe the amount of deviation from or similarity with prototypical activations seen in young adults during novelty processing and successful encoding, by focusing on either typical versus atypical activations (FADE, functional activity deviation during encoding) or activations and deactivations (SAME, similarity of activations during memory encoding). These scores might constitute more robust predictors than voxel-wise fMRI contrasts, as a recent meta-analysis suggested that test-retest reliability of task-based fMRI is mediocre, and the authors recommended whole-brain aggregate analysis rather than voxel- or ROI-based analyses to improve reliability (Elliott et al., 2020).

As an intermediate variable between task-based fMRI and structural MRI, we included the strength of resting-state fMRI signal fluctuations (Jia et al., 2020). Although resting-state fMRI, like task-based fMRI, measures the BOLD signal, it is, like structural MRI, not selective with respect to specific cognitive functions, because subjects are not performing a specific cognitive task (Buckner et al., 2008).

We hypothesized that both chronological age and memory performance could be best predicted from structural MRI, because age-related decrease of memory performance is typically accompanied by structural brain alterations (Cabeza et al., 2004; de Mooij et al., 2018). Whether any MRI modality would outperform the others’ prediction of memory performance, was assessed exploratively.

## 2. Methods

### 2.1. Participants

The study was approved by the Ethics Committee of the Otto von Guericke University Magdeburg, Faculty of Medicine, and written informed consent was obtained from all participants in accordance with the Declaration of Helsinki (World Medical Association, 2013). Participants were recruited via flyers at the local universities (mainly young subjects), advertisements in local newspapers (mainly older participants) and during public outreach events of the institute (e.g., *Long Night of the Sciences*).

The study cohort consisted of a total of 259 neurologically and psychiatrically healthy adults, including 106 young (47 male, 59 female, age range 18-35, mean age 24.12 ± 4.00 years) and 153 older (59 male, 94 female, age range 51-80, mean age 64.04 ± 6.74 years) participants. According to self-report, all participants were right-handed and did not use neurological or psychiatric medication. The Mini-International Neuropsychiatric Interview (M.I.N.I.; Sheehan et al., 1998; German version by Ackenheil et al., 1999) was used to exclude present or past psychiatric illness, alcohol or drug dependence.

Please note that this study is based on the same participant sample as described in Soch et al. (2021a, 2021b) and Richter et al. (2022). The analyses and results described in this study are novel and have not been described or shown elsewhere.

### 2.2. Experimental paradigm

During the fMRI experiment, participants performed a visual memory encoding paradigm with an indoor/outdoor judgment as the incidental encoding task. Compared to earlier publications of this paradigm (Düzel et al., 2011; Barman et al., 2014; Schott et al., 2014; Assmann et al., 2020), the trial timings had been adapted as part of the DZNE-Longitudinal Cognitive Impairment and Dementia (DELCODE) study protocol (Düzel et al., 2018; Bainbridge et al., 2019; see Soch et al., 2021a, for a detailed comparison of trial timings and acquisition parameters). Subjects viewed photographs showing indoor and outdoor scenes, which were either novel at the time of presentation (44 indoor and 44 outdoor scenes) or were repetitions of two highly familiar “master” images (22 indoor and 22 outdoor trials), i.e. one indoor and one outdoor scene pre-familiarized before the actual experiment (cf. Soch et al., 2021a, Fig. 1B). Thus, every subject was presented with 88 unique images and 2 master images that were presented 22 times each. Participants were instructed to categorize images as “indoor” or “outdoor” via button press. Each picture was presented for 2.5 s, followed by a variable delay between 0.70 s and 2.65 s. To optimize estimation of the condition-specific BOLD responses despite the short delay, simulations were employed to optimize the trial order and jitter, as described previously (Hinrichs et al., 2000; Düzel et al., 2011).

**Figure 1.**
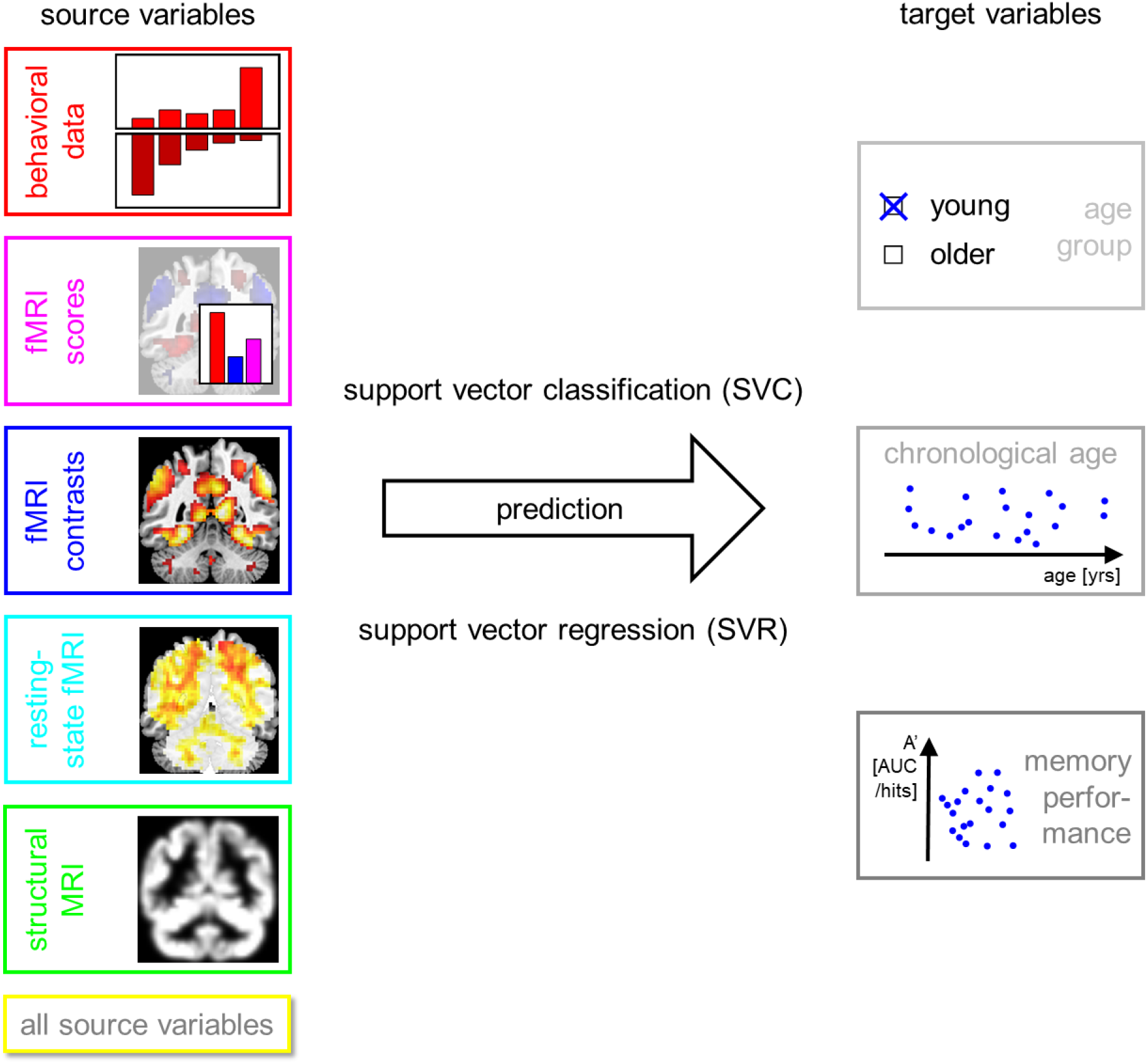
Methodology of the present study. Several target variables of interest (right) are predicted using several sets of source variables (left), thought to be markers of cognitive decline in old age, using machine learning techniques (center).

Approximately 70 minutes (70.23 ± 3.77 min) after the start of the fMRI session, subjects performed a computer-based recognition memory test outside the scanner, in which they were presented with the 88 images that were shown once during the fMRI encoding phase (*old*) and 44 images they had not seen before (*new*). Participants rated each image on a five-point Likert scale from 1 (“definitely new”) to 5 (“definitely old”). For detailed experimental procedure, see Assmann et al. (2020) and Soch et al. (2021a).

### 2.3. fMRI data acquisition

Structural and functional MRI data were acquired on two Siemens 3T MR tomographs (Siemens Verio: 58 young, 83 older; Siemens Skyra: 48 young, 70 older), following the exact same protocol used in the DELCODE study (Jessen et al., 2018; Düzel et al., 2019; Billete et al., in review).

A T1-weighted MPRAGE image (TR = 2.5 s, TE = 4.37 ms, flip-α = 7°; 192 slices, 256 × 256 in-plane resolution, voxel size = 1 × 1 × 1 mm) was acquired for co-registration and improved spatial normalization. Phase and magnitude fieldmap images were acquired to improve correction for artifacts resulting from magnetic field inhomogeneities (see below).

For functional MRI (fMRI), 206 T2*-weighted echo-planar images (EPIs; TR = 2.58 s, TE = 30 ms, flip-α = 80°; 47 slices, 64 × 64 in-plane resolution, voxel size = 3.5 × 3.5 × 3.5 mm) were acquired in interleaved-ascending slice order (1, 3, ..47, 2, 4, ..46). Prior to this task-based fMRI experiment, a resting-state fMRI run was acquired, comprising 180 EPIs with otherwise identical acquisition parameters. The total scanning times were 531.48 s (≈ 9:51 min) for the task-based fMRI run and 464.4 s (≈ 7:44 min) for the resting-state fMRI session. The complete study protocol also included a T2-weighted MR image in perpendicular orientation to the hippocampal axis (TR = 3.5 s, TE = 350 ms, 64 slices, voxel size = 0.5 × 0.5 × 1.5 mm) for optimized segmentation of the hippocampus (Dounavi et al., 2020) and additional structural imaging not used in the analyses reported here.

### 2.4. fMRI data preprocessing

Data preprocessing was performed using Statistical Parametric Mapping (SPM12; Wellcome Trust Center for Neuroimaging, University College London, London, UK). EPIs were corrected for acquisition time delay (*slice timing*), head motion (*realignment*) and magnetic field inhomogeneities (*unwarping*), using voxel-displacement maps (VDMs) derived from the fieldmaps. The MPRAGE image was spatially co-registered to the mean unwarped image and *segmented* into six tissue types, using the unified segmentation and normalization algorithm implemented in SPM12. The resulting forward deformation parameters were used to *normalize* unwarped EPIs into a standard stereotactic reference frame (Montreal Neurological Institute, MNI; voxel size = 3 × 3 × 3 mm). Normalized images were spatially *smoothed* using an isotropic Gaussian kernel of 6 mm full width at half maximum (FWHM).

### 2.5. General linear modelling

For first-level fMRI data analysis, which was also performed in SPM12, we used a parametric general linear model (GLM) of the subsequent memory effect that has recently been demonstrated to outperform the so far more commonly employed categorical models of fMRI subsequent memory effects (Soch et al., 2021a) when subsequent memory responses are recorded as memory confidence ratings on a parametric scale.

This model included two onset regressors, one for novel images at the time of presentation (“novelty regressor”) and one for presentations of the two pre-familiarized images (“master regressor”). Both regressors were created as short box-car stimulus functions with an event duration of 2.5 s, convolved with the canonical hemodynamic response function, as implemented in SPM12.

The regressor reflecting subsequent memory performance was obtained by parametrically modulating the novelty regressor with a function describing subsequent memory report. Specifically, the parametric modulator (PM) was given by

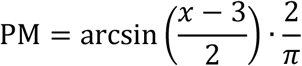

where *x* ∈ {1, 2, 3,4, 5}is the subsequent memory report, such that - 1 ≤PM≤+1. Compared to a linear-parametric model, this transformation puts a higher weight on definitely remembered (5) or forgotten (1) items compared to probably remembered (4) or forgotten (2) items (Soch et al., 2021a, Fig. 2A).

**Figure 2.**
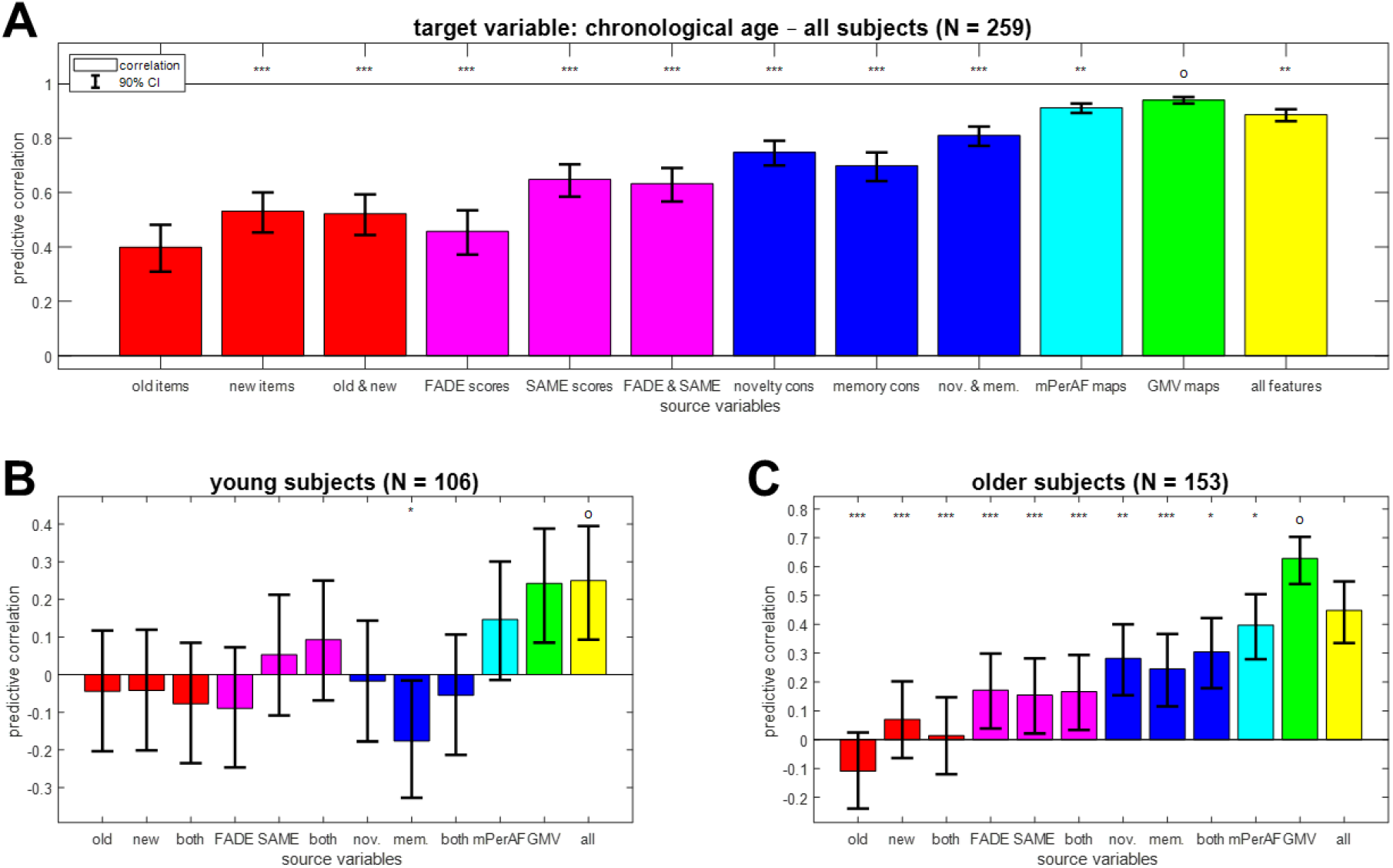
Prediction of chronological age from different feature sets. Bar plots show correlation coefficients for predicting chronological age (in years) **(A)** across all subjects, **(B)** in young subjects only or **(C)** in older subjects only from behavioral data (red), fMRI scores (magenta), task-based fMRI contrasts (blue), resting-state fMRI maps (cyan) and structural MRI (green), or all features (yellow). Error bars denote 90% confidence intervals; x-axis labels are explained in Table 4. The feature set with the highest predictive correlation is denoted with an “o”; other feature sets are labeled with asterisks to indicate significantly different mean absolute error (* p < 0.05, ** p < 0.01, *** p < 0.001, otherwise not significant).

The model also included the six rigid-body movement parameters obtained from realignment as covariates of no interest and a constant representing the implicit baseline.

### 2.6. Extraction of target variables

For each subject, age group (young vs. older), chronological age (in years) and memory performance (area under the curve, AUC; see Soch et al., 2021b, Appendix B) were extracted as dependent variables, i.e. target variables for prediction analyses (see Table 1).

**Table 1.**
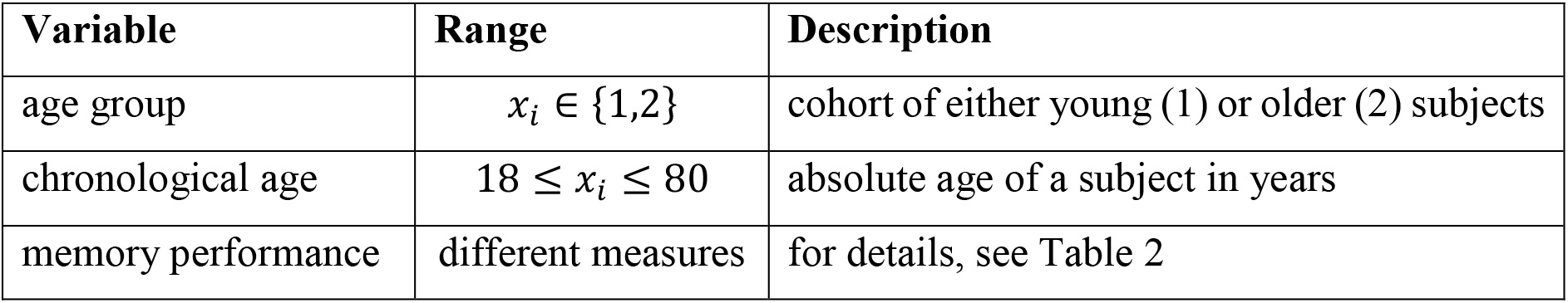
Target variables used for prediction analyses. Details on the different measures of memory performance are given in Table 2.

Note that our measure of memory performance is not completely independent from some of the source variables, because it was obtained from the same task during which behavioral data and functional MRI were acquired (see Section 2.7). For this reason, we also used independent measures of memory performance to test the predictive performance of our candidate variables. These measures include (i) the number of items retrieved in a verbal learning task (verbal learning and memory test, VLMT; Helmstaedter et al., 2001), in a recall after 30 minutes or 1 day; and (ii) the number of points obtained in a semantic memory test (Wechsler memory scale, WMS; Härting et al., 2000), in a recall after 30 minutes or 1 day (see Table 2). For detailed description of these neuropsychological assessments, see Richter et al. (2022).

**Table 2.** Measures of memory performance used as target variables. Abbreviations: FADE = name of the fMRI paradigm; A’ = area under the curve (AUC) when plotting the hit rate as a function of false alarm rate; VLMT = verbal learning and memory test; WMS = Wechsler memory scale.

| Measure | Stimulus material | Encoding type | Recall delay | Recall type | Theoretical range | Actual range |
| --- | --- | --- | --- | --- | --- | --- |
| FADE-A' | visual | incidental | 70 min | recognition | 0 – 1 | 0.53 – 0.98 |
| VLMT-30min | verbal | intentional | 30 min | free recall | 0 – 15 | 4 – 15 |
| VLMT-1d | verbal | intentional | 1 d | free recall | 0 – 15 | 2 – 15 |
| WMS-30min | auditory | intentional | 30 min | free recall | 0 – 50 | 9 – 46 |
| WMS-1d | auditory | intentional | 1d | free recall | 0 – 50 | 6 – 45 |

### 2.7. Extraction of source variables

For each subject, the following variables were extracted as independent variables, i.e. source variables for prediction analyses (see Table 3):

- *behavioral response frequencies:* In the surprise recognition memory test, subjects provided memory confidence ratings between 1 and 5 for all 88 old stimuli, (i.e. items presented during the encoding session) and 44 new stimuli (i.e. items not seen during the encoding session; see Section 2.2). From the responses of subject *i*, we calculated *O_ij_*, the proportion of old items rated with confidence level *j*, and *n_ij_*, the proportion of new items rated with *j*. The variables *o*_*i*3_ and *n*_*i*3_ were dropped to avoid collinearity of predictor variables, since all “old” proportions and all “new” proportions added up to 1, respectively.
- *fMRI contrast images:* The GLM for first-level fMRI data analysis contained one regressor for novel images, parametrically modulated with a non-linear transformation of memory confidence, and another regressor for master images (see Section 2.5). From this, we generated fMRI contrast maps for “novelty processing” as such, by subtracting the master regressor from the novelty regressor, and for “subsequent memory” effects, identical to the estimated regression coefficient for the parametric modulator.
- *fMRI summary statistics:* We then identified regions with group-level significant positive and negative activations on these contrasts in young subjects. Using these voxels as masks, we calculated two recently described fMRI scores quantifying the deviation of older adults from the prototypical activation of young subjects (for detailed procedure and extracted scores, see Soch et al., 2021b, Sections 2.6 to 2.8). Both scores, FADE-classic (FADE = functional activity deviation during encoding; Düzel et al., 2011) and FADE-SAME (SAME =similarities of activations during memory encoding; Soch et al., 2021b), were computed from both contrasts, novelty processing and subsequent memory.
- *resting-state fMRI maps:* We then applied the RESTplus toolbox (Jia et al., 2019) to the preprocessed resting-state fMRI scans of each subject and calculated the voxel-wise percent of amplitude fluctuation (PerAF) of signals in the frequency range from 0.01 to 0.08 Hz. PerAF is the average absolute deviation from the signal mean, measured in percent (Jia et al., 2020, eq. 1). Here, we used “mean PerAF” (mPerAF), which additionally divides PerAF by the global mean (Jia et al., 2020, Tab. 1) and was already employed in a previous study (Kizilirmak et al., in prep.).
- *structural MRI maps:* Finally, the T1 image of each subject was submitted to structural MRI analyses (i.e. voxel-based morphometry, VBM) using the Computational Anatomy Toolbox (CAT12; Structural Brain Mapping Group, Department of Neurology, University Jena, Germany), resulting in gray matter volume (GMV) maps. These maps were additionally smoothed using a Gaussian kernel (isotropic FWHM = 6 mm) before entering whole-brain decoding analyses.

**Table 3.** Source variables used for prediction analyses. Abbreviations: FADE = functional activity deviation during encoding, SAME = similarities of activations during memory encoding, 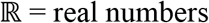, *ν* = number of (in-mask) voxels.

| Variables | Range | Description |
| --- | --- | --- |
| behavioral<br>response<br>frequencies | $o_{i1}, \dots, o_{i5} \in [0,1]$<br>$n_{i1}, \dots, n_{i5} \in [0,1]$ | proportion of old items replied to with 1, ..., 5 and<br>proportion of new items replied to with 1, ..., 5 |
| fMRI<br>summary<br>statistics | $y_{i1}, \dots, y_{i4} \in \mathbb{R}$ | two scores (FADE-classic, FADE-SAME) computed<br>from two fMRI contrasts (novelty processing,<br>subsequent memory) |
| fMRI<br>contrast<br>images | $Y_i \in \mathbb{R}^v$ | voxel-wise fMRI contrasts computed in SPM,<br>representing activations related to novelty processing<br>(novel images – master images) or subsequent me-<br>mory (parametric modulator with memory response) |
| resting-state<br>fMRI maps | $Y_i \in \mathbb{R}^v$ | voxel-wise percent of amplitude fluctuation (mPerAF)<br>computed using the REST toolbox, based on fMRI<br>signals measured during a resting-state session |
| structural<br>MRI maps | $Y_i \in \mathbb{R}^v$ | voxel-wise gray matter volumes computed in CAT12,<br>based on each subject's T1 image |

### 2.8. Prediction of target from source variables

After source and target variables were extracted, several analyses were performed and each analysis consisted in predicting a single target variable from a feature set of source variables using support vector machines (SVM; see Figure 1 and Table 4).

**Table 4.**
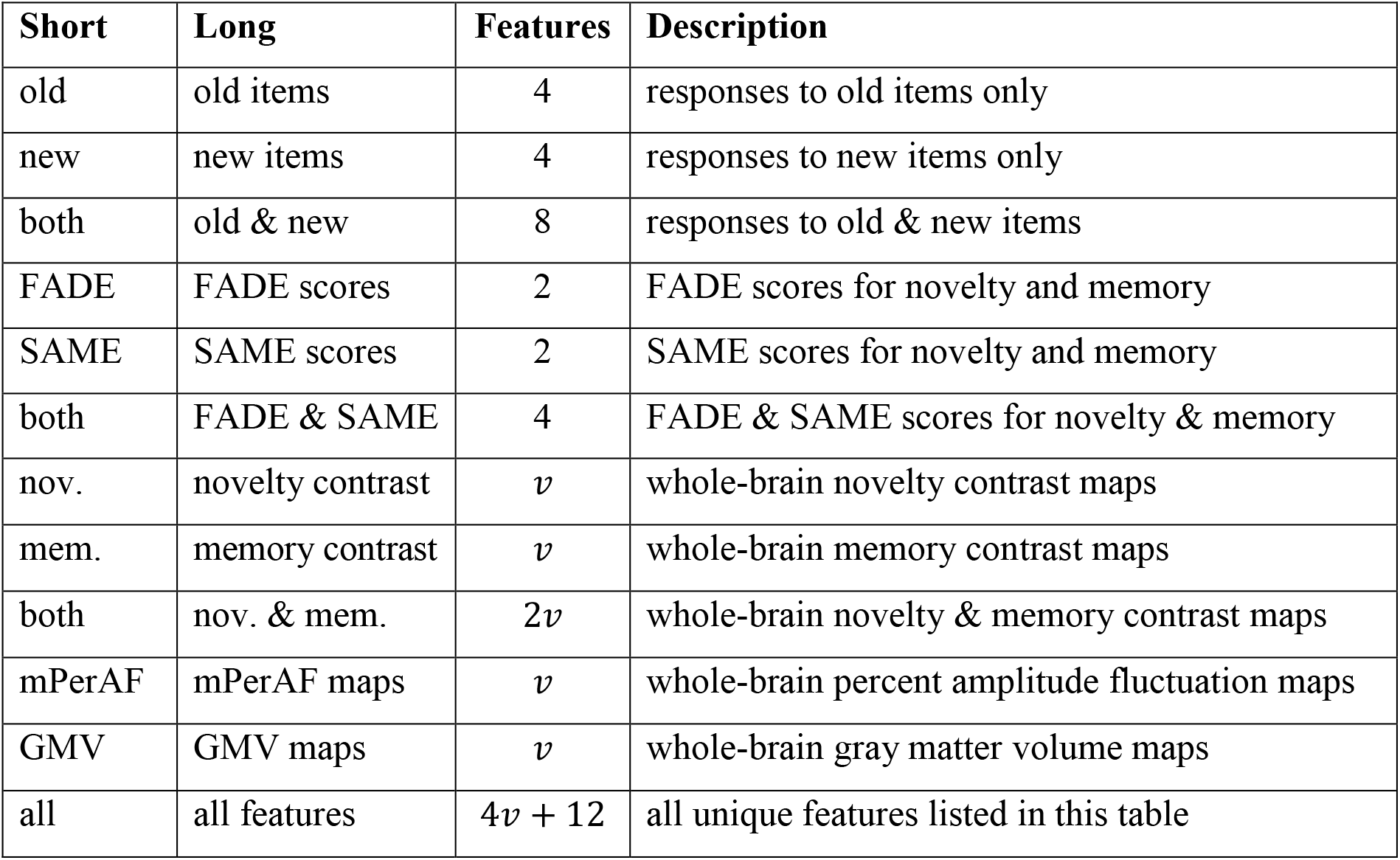
Feature sets used for prediction analyses. Short and long feature set names are used as x-axis labels on Figures 2–5. The number of features corresponds to the number of columns in the data matrix used for prediction. Abbreviations: FADE = functional activity deviation during encoding, SAME = similarities of activations during memory encoding, *ν* = number of (in-mask) voxels.

For decoding the age group a subject was belonging to, we used support vector classification (SVC) using a linear SVM with C = 1. For predicting chronological age and memory performance, we used support vector regression (SVR) using a linear SVM with C = 1. For both, SVC and SVR, subjects were split with k-fold cross-validation (CV) on subjects per group using k = 10 CV folds. All SVM analyses were implemented using LibSVM in MATLAB via in-house scripts available from GitHub^1^.

### 2.9. Distributional transformation

When predicting chronological age and memory performance, distributional transformation (DT) was applied to preserve the observed distribution of the target variable (Soch, 2020). DT is a post-processing operation that maps predicted values to the variable’s distribution in the training data and can improve prediction precision.

For example, memory measured as AUC always falls into the range between 0 and 1, but a trained SVM may also return values smaller than 0 or larger than 1. Then, DT brings predicted values into the natural range of the target variable while keeping the ranks of all predicted values identical before and after transformation (Soch, 2020). The same holds when predicting age which was always between 18 and 80 years in our study. For subgroup analyses, only the age range of the respective group (young vs. older) was applied.

### 2.10. Performance assessment

The prediction precision was assessed using balanced accuracy (ranging between 0 and 1) when decoding age group, i.e. by averaging the decoding accuracies for young and older subjects (Brodersen et al., 2010); and using correlation coefficients (ranging between –1 and +1) when predicting chronological age and memory performance, i.e. as the sample correlation coefficient between actual and predicted values of those variables. For each precision measure, a 90% confidence interval was established.^2^

When predicting chronological age and memory performance, we additionally calculated absolute errors (AE) between predicted and actual target values and submitted them to Wilcoxon signed-rank tests to check for significant reduction of the mean absolute error (MAE) from one feature set to another. This non-parametric test was chosen due to the presumably non-normal distribution of absolute errors. For each target variable, AEs of the feature set with the highest correlation coefficient were compared against AEs of each other feature set to test whether performances of the feature sets were significantly different from that of the most predictive feature set (see e.g. Figure 3).

**Figure 3.**
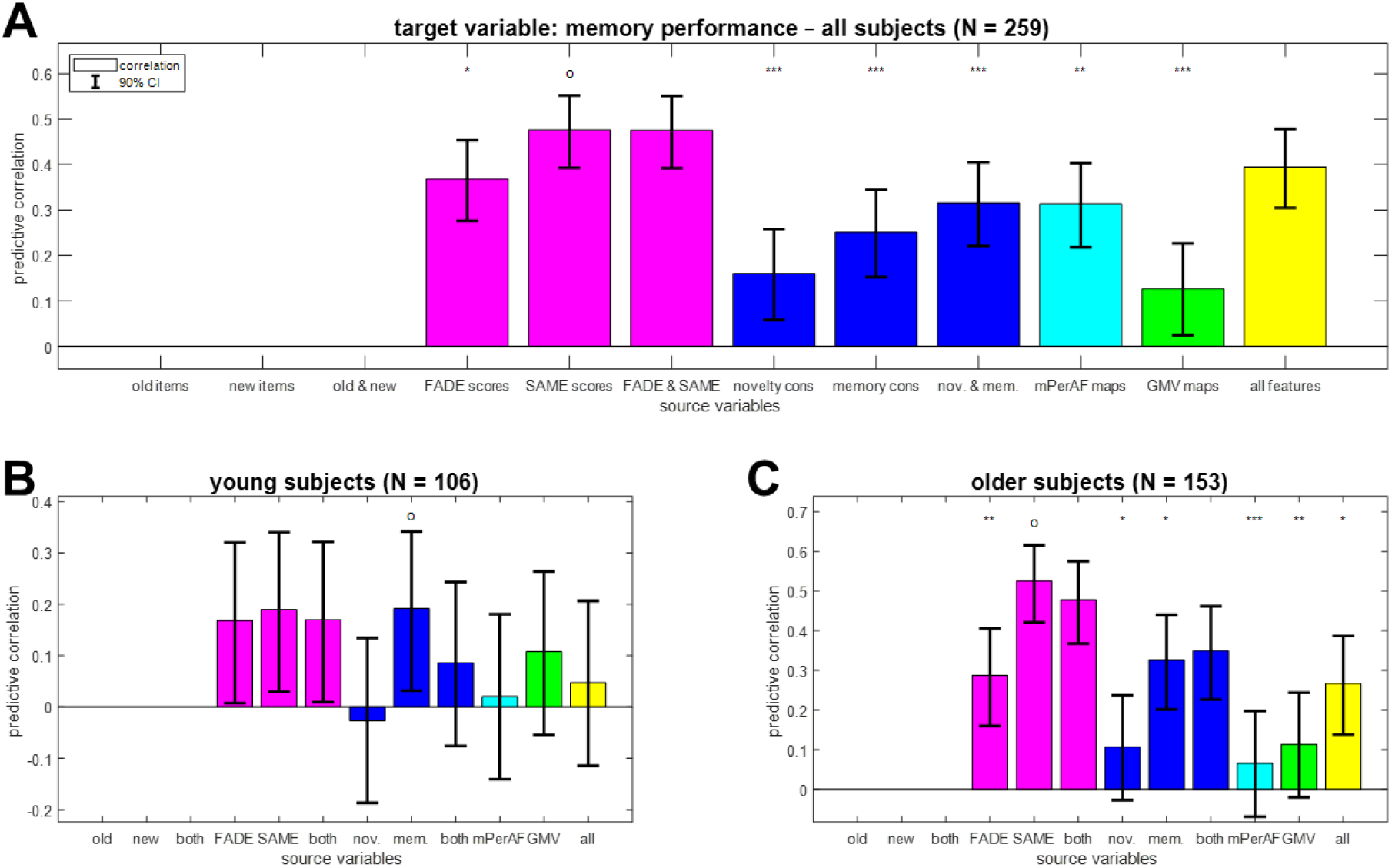
Reconstruction of memory performance from different feature sets. Bar plots show correlation coefficients for predicting memory performance (area under the curve) **(A)** across all subjects, **(B)** in young subjects only or **(C)** in older subjects only from fMRI scores (magenta), task-based fMRI contrasts (blue), resting-state fMRI maps (cyan) and structural MRI (green), or all features (yellow). Note that memory performance can be directly derived from behavioral data which is why the corresponding prediction analyses were not performed. The layout follows that of Figure 2.

## 3. Results

### 3.1. Chronological age is best predicted from structural MRI maps

The age group a subject belonged to (young vs. older subjects) could be predicted from all feature sets with above-chance decoding accuracy (see Supplementary Figure S1). The highest accuracy was obtained with GMV maps (balanced accuracy, BA = 96.01 %; confidence interval, CI = [0.931, 0.976]) and the lowest accuracy was obtained with response frequencies to old items (BA = 59.68 %, CI = [0.542, 0.646]).

When predicting chronological age (in years) across all subjects, we found significant correlations for all feature sets (see Figure 2A; old items: r = 0.40; GMV maps: r = 0.95). However, this was mainly attributable to the inherent correlation between chronological age and age group (see Section 2.1), such that decoding age group is already a good predictor for chronological age. Therefore, we performed the same analyses separately within young subjects (18-35 years) and within older subjects (60-80 years).

In young subjects, chronological age could only be reconstructed from whole-brain GMV maps (see Figure 2B; r = 0.24, CI = [0.085, 0.388]; all other |r| < 0.20). In older subjects, chronological age could be predicted from GMV and resting-state fMRI maps (see Figure 2C; GMV maps: r = 0.63, CI = [0.540, 0.703]; mPerAF maps: r = 0.40, CI = [0.279, 0.504]) and, with lower accuracy, from task-based fMRI contrasts (novelty & memory: r = 0.30, CI = [0.179, 0.421]) and fMRI summary statistics (FADE & SAME: r = 0.17, CI = [0.033, 0.293]), but not from behavioral response frequencies (old & new: r = 0.01, CI = [-0.120, 0.147]).

### 3.2. Dependent memory performance is best predicted from task-based fMRI

Similar to chronological age, memory performance (AUC) across all subjects could be predicted from all feature sets^3^ (see Figure 3A; GMV maps: r = 0.13; SAME scores: r = 0.48). However, as memory performance is also strongly influenced by age group, with young subjects performing significantly better than older subjects (young: μ1 = 0.82; older: μ2 = 0.77; effect size: d’= 0.72; two-sample t-test: t = 5.67, p < 0.001), we again analyzed this target variable separately within young and older subjects, respectively.

In both age groups, memory performance predicted by GMV maps was not correlated to actual memory performance (young: r = 0.11; older: r = 0.11). Instead, memory performance was best predicted by the fMRI memory contrast in young subjects (see Figure 3B; r = 0.19, CI = [0.032, 0.342]) and the SAME scores in older subjects (see Figure 3C; r = 0.53, CI = [0.421, 0.616]). Note that the predictive accuracy when predicting from just four single-value fMRI scores (FADE & SAME: r = 0.48, CI = [0.368, 0.575]) was better than using two whole-brain task-based fMRI contrasts (novelty & memory: r = 0.35, CI = [0.227, 0.461]).

### 3.3. Independent memory performance is best predicted from single-value fMRI scores

When predicting independent measures of memory performance (see Section 2.6 and Table 2), we restrict the results report to the older subjects, because those measures could not be reliably predicted at all in young subjects (see Supplementary Figure S2), probably due to the lower variation in their close-to-ceiling memory performance.

Generally, the prediction of memory performance in independent tests was less accurate than that of behavioral memory performance in the fMRI task itself (cf. Figure 4 vs. 3C). Besides this, outcomes from all memory tests are best predicted by the SAME scores (see Figure 4A/B/D; VLMT 30 min: r = 0.25, CI = [0.124, 0.375]; VLMT 1d: r = 0.23, CI = [0.099, 0.356]; WMS 1d: r = 0.33, CI = [0.198, 0.442]) or FADE scores (see Figure 4C; WMS 30 min: r = 0.36, CI = [0.234, 0.471]).

**Figure 4.**
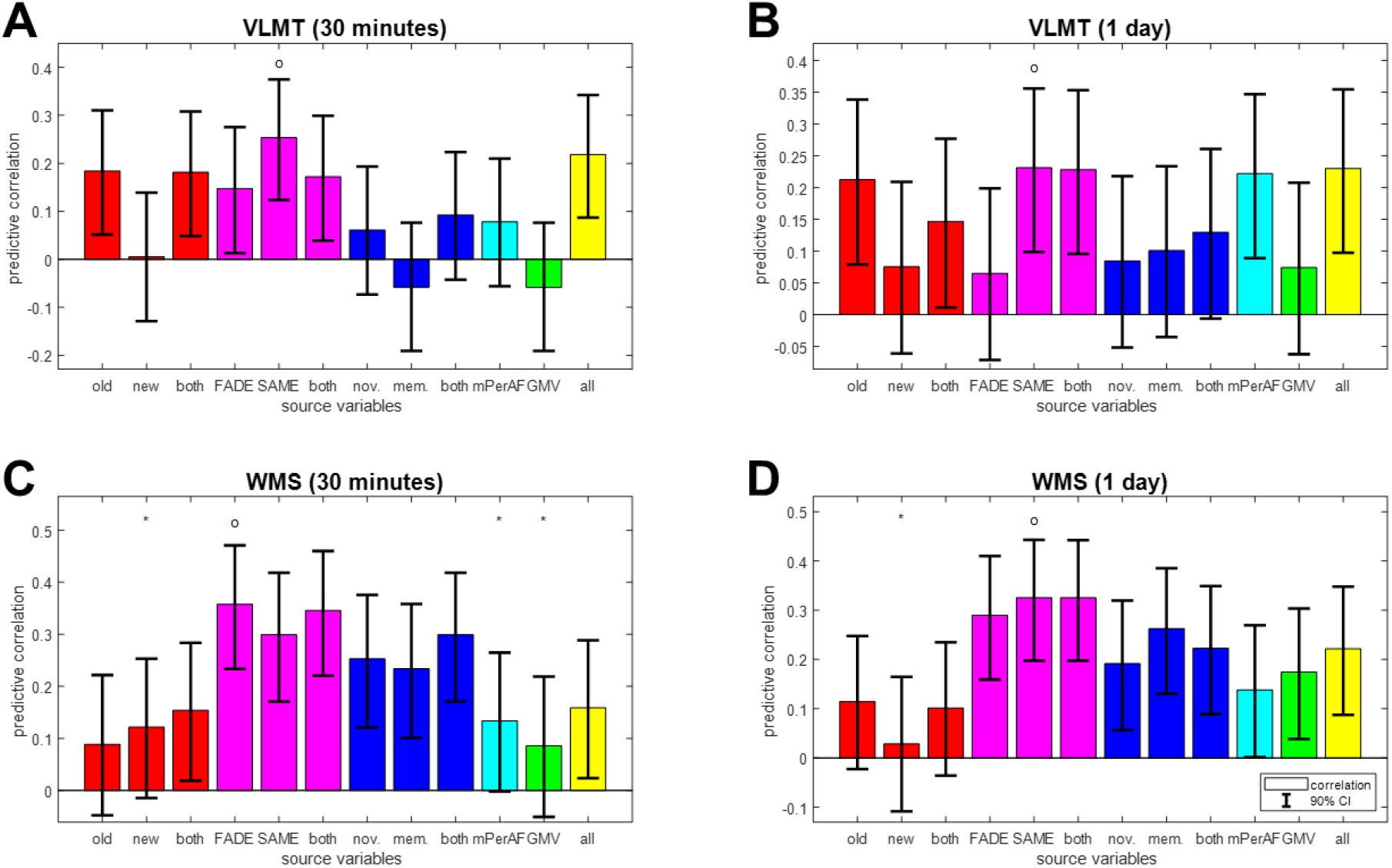
Reconstruction of independent memory performance in older subjects. Bar plots show correlation coefficients for predicting, in older subjects only, independent measures of memory performance, namely **(A)** VLMT items after 30 minutes, **(B)** VLMT items after 1 day, **(C)** WMS points after 30 minutes and **(D)** WMS points after 1 day, from behavioral data (red), fMRI scores (magenta), task-based fMRI contrasts (blue), resting-state fMRI maps (cyan) and structural MRI (green), or all features (yellow). The layout follows that of Figure 3C.

Moreover, there appears to be a dissociation by type of memory test: Whereas performance in the verbal-semantic VLMT could be predicted from behavioral responses to old items, but not task-based fMRI contrast maps, the reverse pattern was seen for performance in the auditory-episodic WMS (see Figure 4, red and blue bars; see Supplementary Discussion for potential explanations). Notably, the two SAME scores and all four fMRI-based scores were the only feature sets that allowed for above-chance prediction of all four independent measures of memory performance (see Figure 4, magenta bars).

### 3.4. Effects of age and memory are specific to structural vs. functional MRI

To follow up on the findings of predictive analyses, especially the differences in predicting participants’ age vs. memory (cf. Figure 2C vs. 3C), we explicitly compared functional and structural MRI data in older subjects using sub-group analyses. To this end, we partitioned all older subjects into four groups based on (i) chronological age, separating into “young” and “old” older subjects; and (ii) memory performance, separating higher from lower memory performance subjects (see Supplementary Figure S3). Then, the voxel-wise data of the quarter with the lowest values and the quarter with the highest values were submitted to second-level two-sample t-tests in SPM. This analysis was performed for both fMRI contrasts, mPerAF maps and GMV maps. Thresholded statistical parametric maps were FWE-cluster-corrected (cluster-defining threshold, CDT: p < 0.001, k = 0), resulting in a minimum cluster size for each analysis (novelty: k = 42; memory: k = 27; mPerAF: k = 23; GMV: k = 33 (separating by age) and k = 42 (separating by memory); see Figure 5).

**Figure 5.**
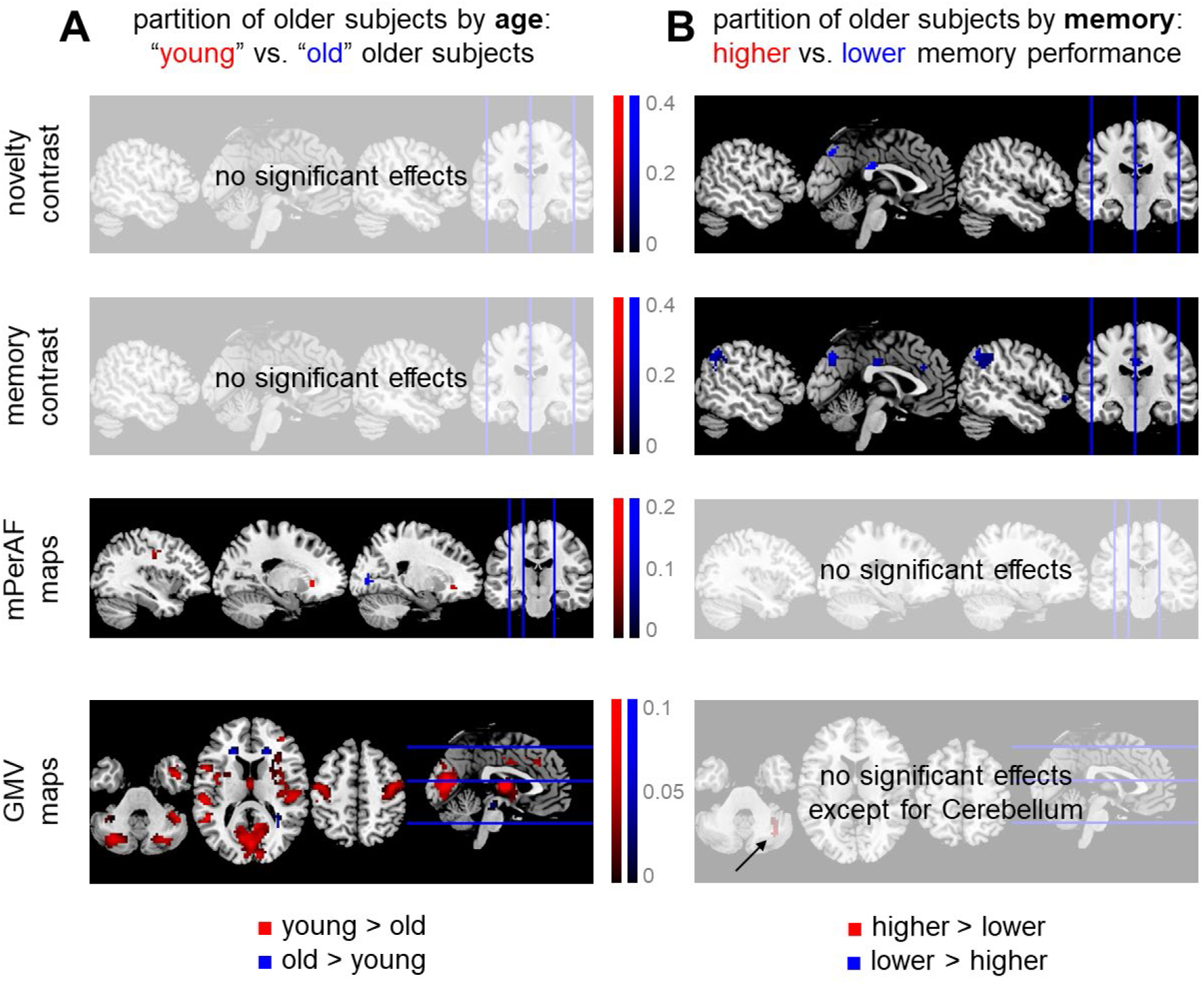
Differential effects of age and memory in structural and functional MRI. Significant differences **(A)** between “young” and “old” older subjects and **(B)** between older subjects with higher vs. lower memory performance, with respect to fMRI activity during novelty processing (1st row), subsequent memory (2nd row), fMRI amplitudes during rest (3rd row) and voxel-wise gray matter volume (4th row). Thresholded SPMs are FWE-corrected for cluster size (CDT: p < 0.001, k = 0). Colored voxels indicate significantly higher values for either young subjects and those with higher memory performance (red) or old subjects and those with lower memory performance (blue).

Taken together, we observed a double dissociation of structural MRI versus task-based fMRI and age versus memory, in the sense that (i) when partitioning subjects by chronological age, there were significant effects on structural MRI (see Figure 5A); and (ii) when partitioning subjects by memory performance, there were significant effects on task-based fMRI (see Figure 5B); at the same time, there were no age-related differences with respect to task-based fMRI and no memory-related differences with respect to structural MRI. Resting-state fMRI maps showed differences between younger and older subjects, but not between those with high vs. low memory performance (see Figure 5, 3rd row), suggesting that their informational content is closer to structural MRI than to task-based fMRI.

### 3.5. Single-value fMRI scores have moderate predictive utility

To assess the predictive utility of fMRI summary statistics, we used FADE and SAME scores computed from novelty and memory contrasts (i.e. four features, cf. Table 4) and evaluated the precision by which these scores predict memory performance in two ways.

First, we compared predicted with actual values when reconstructing area under the curve (AUC) in the fMRI memory paradigm from FADE and SAME scores (cf. Figure 3B/C). In older subjects, there was a correlation of 0.47 (p < 0.001) and AUC could be predicted with a mean absolute error (MAE) of 0.06 (see Figure 6B). For comparison, the same correlation was 0.17 (p = 0.082) with an MAE of 0.08 in young subjects (see Figure 6A).

Second, we tested how well sub-groups of the older subjects formed for the previous analysis (see Section 3.4 and cf. Figure 5A/B) could be classified from fMRI scores. When classifying older subjects with lower vs. higher memory performance based on FADE and SAME scores (N = 76), the decoding accuracy was 72.37 % (sensitivity: 76.32 %; specificity: 68.42 %). For comparison, the decoding accuracy was 84.93 % (sensitivity: 81.08 %; specificity: 88.89 %) when classifying “old” vs. “young” older subjects based on GMV maps (N = 73).

**Figure 6.**
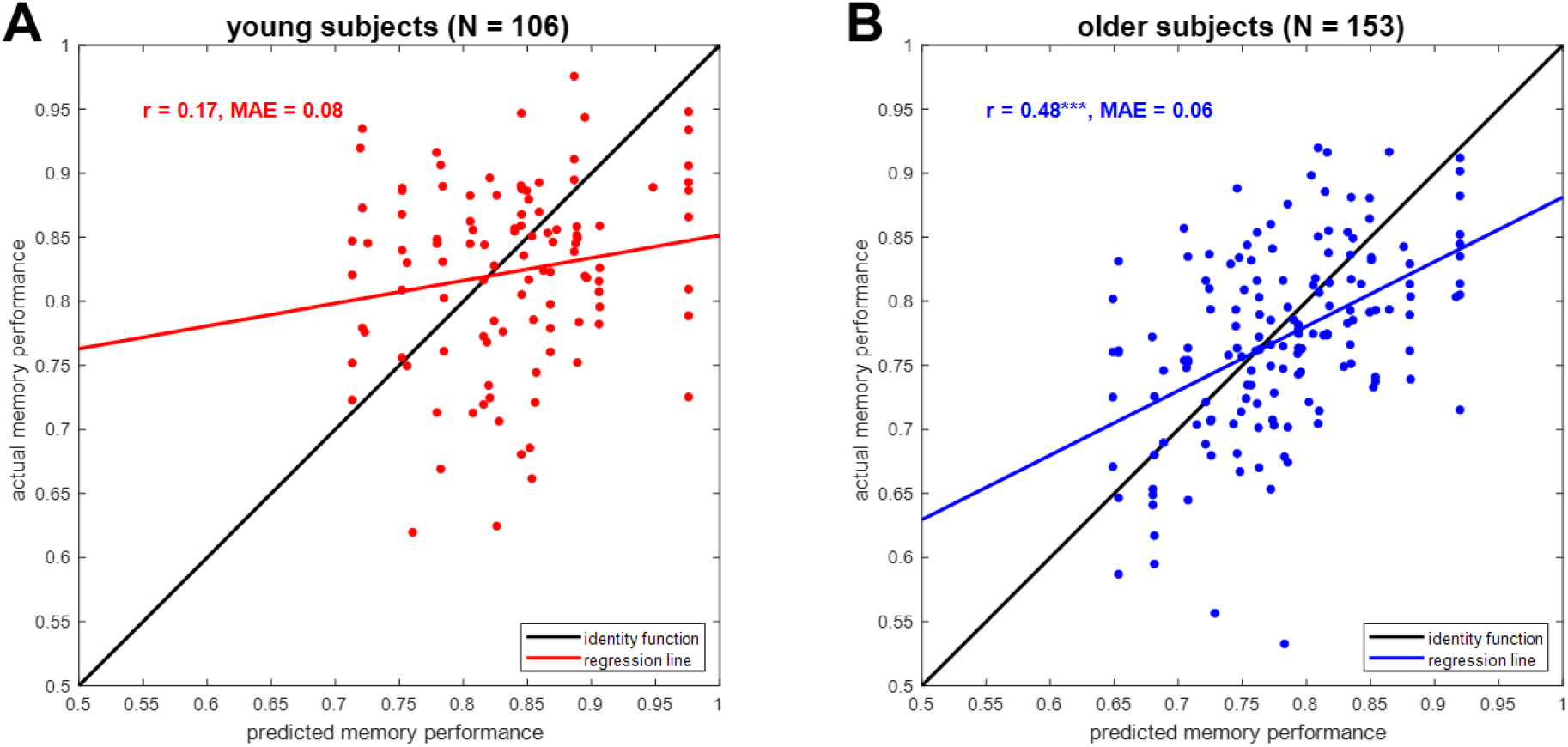
Prediction of memory performance from single-value fMRI scores. Scatter plots of actual vs. predicted memory performance when reconstructing memory performance from FADE and SAME scores (see Figure 3, magenta bars) in **(A)** young subjects and **(B)** older subjects. Abbreviations: r = correlation coefficient, MAE = mean absolute error, *** p < 0.001.

## 4. Discussion

In the present study, we have comparatively evaluated the ability of structural and functional (resting-state and task-based) MRI data as well as behavioral measures to predict chronological age versus memory performance in young and older healthy adults (see Figure 1). While all modalities could predict age group, within-group prediction of age and memory performance revealed distinct patterns. Among young and older subjects, chronological age was best predicted by structural MRI and also resting-state fMRI (see Figure 2B and 2C), whereas memory performance was best predicted by functional MRI contrasts (novelty and subsequent memory effects) and especially single-value fMRI-based scores (see Figure 3C and 4) in older participants only.

### 4.1. Prediction of chronological age from structural MRI

All of the candidate predictors employed in the present study have previously been shown to exhibit age-related differences: (i) behavioral memory responses are different between age groups, with older adults producing more false positives which reduces memory performance (cf. Soch et al., 2021a, Tab. S2; also see Duarte et al., 2010); (ii) memory-related fMRI responses differ between age groups, with older adults showing reduced parahippocampal activations and reduced default mode network (DMN) deactivations during novelty processing and subsequent memory (cf. Soch et al., 2021b, Fig. 2; also see Maillet and Rajah, 2014; Billette et al., in review); (iii) resting-state fMRI patterns exhibit global age-related differences (Foo et al., 2021; Xing, 2021), and (iv) quantitative structural MRI approaches like VBM yield robust and well-replicated age-related differences, with older adults showing reduced hippocampal volumes (cf. Kizilirmak et al., in prep., Fig. 3A; also see Veldsman et al., 2020) as well as reduced cortical and subcortical GMV, particularly in structures of the human memory network like the medial temporal lobe (Schiltz et al., 2006; Minkova et al., 2017).

In line with the aforementioned observations, all variables could discriminate between age groups, but within the group of older adults, a distinct pattern emerged regarding the prediction of chronological age and memory performance, respectively. Chronological age was best predicted from voxel-wise GMV, reflecting the well-replicated observation that both cortical and subcortical GM show age-related volume loss (Minkova et al., 2017; Soch, 2020; Veldsman et al., 2020), which is, longitudinally, already observable within a year’s time (Fjell et al., 2009, 2013; Bagarinao et al., 2022). Predictive correlation of whole-brain GMV and chronological age within the group of older adults was, however, only moderate, most likely reflecting the considerable inter-individual variability in age-related structural brain changes. This phenomenon has in fact been conceptualized within the brain-age framework, a widely researched approach to employ differences between predicted brain age and chronological age as a biomarker for brain health in aging (see, e.g., Cole and Franke, 2017; Bashyam et al., 2020). Including other predictors in the model did not improve age prediction among older adults (Figure 2C), suggesting that the biological information actually predicting chronological rather than brain age might be limited.

In a recent competition to predict chronological age from structural neuroimaging (Fisch et al., 2020), the winning performance, a mean absolute error of 2.90 years, was achieved using lightweight 3D convolutional neural networks (Gong et al., 2020). Moreover, it was shown that distributional transformation can improve the MAE by about half a year, utilizing the distribution of the target values in the training data (Soch, 2020), an approach that was also used in the present study (see Section 2.9).

### 4.2. Functional MRI as predictor of cognitive performance in old age

Unlike chronological age, memory performance could not be reliably predicted from GMV. This is compatible with the fact that in previous studies, we found no correlations between hippocampal volume and our task-based fMRI summary statistics for both hemispheres, using two scores, computed from two contrasts (Soch et al., 2021b, Fig. 4). It is also supported by another study, in which a combination of ApoE genotype and task-based fMRI was identified as the best predictor of cognitive decline in healthy older adults (Woodard et al., 2010). In line with those findings, we here observed that memory performance could be predicted from single-value fMRI scores (see Figure 4), especially when extracting both FADE and SAME scores, from both novelty and memory contrasts (Soch et al., 2021b).

It should be noted that the cognitive task underlying our fMRI data set (incidental encoding of visual scenes) in fact targeted declarative long-term memory. In so far, the high predictive value of functional measures derived from activity during such a task (i.e., fMRI novelty and memory contrast maps, FADE and SAME scores) for other measures of declarative memory appears to be a natural outcome, as it is more specifically targeting the to-be-predicted variable than GMV or mPerAF. The same is true for the study of Woodard and colleagues, in which participants encoded names (famous vs. unfamiliar names) and the independent measures of cognitive decline comprised different types of neuropsychological memory assessments. On the other hand, we could recently show that, while the scores derived from the novelty contrast were rather specifically associated with tests of explicit memory, the scores computed from the memory contrast were also associated with measures of global cognition (Richter et al., 2022). More generally, our findings are in line with the notion that cognitive reserve may to a certain degree be independent from structural age-related changes of the brain (Nyberg et al., 2012).

### 4.3. Informational content of resting-state maps

It is also noteworthy that resting-state fMRI behaved more similar to structural MRI than task-based fMRI, with balanced accuracy for mPerAF maps being close to that of GMV maps (see Figure S1) and mPerAF similarly predicting chronological age (see Figure 2C), but not capturing memory performance in older subjects (see Figure 3C). This suggests that at least voxel-wise mPerAF maps derived from resting-state fMRI provide information that is closer to the brain-anatomical information of structural MRI maps than to the neural-processing information of task-based fMRI contrasts.

This is compatible with the line of thought discussed above: While task-based fMRI measures provide informational value for cognitive performance measures, especially when the fMRI task falls into the same cognitive domain as the to-be-predicted performance indicator, resting-state fMRI measures appear to reflect brain integrity more generally (e.g. Mevel et al., 2011).

### 4.4. Successful aging, brain structural integrity, and memory performance

Overall, our results suggest that successful aging, that is, relatively preserved memory in healthy older adults, may not be primarily attributable to lower gray matter loss, but rather to better preserved functional brain networks, as evident in a higher similarity of memory-related brain activity with that of young adults (see Figure 5). This might be different in pathological aging when brain anatomy is affected to a larger extent, but is compatible with earlier studies suggesting that in healthy older adults, functional neurocognitive resources may be more important for cognitive performance than structural measures of brain integrity (Scarmeas et al., 2003; Stern, 2009, 2012; Cabeza et al., 2018).

The observation that structural MRI had no predictive power for memory performance in our study may at first seem surprising, given that there are very large differences with respect to GMV between young and older adults (Farokhian et al., 2017) who typically also differ with respect to memory performance (Soch et al., 2021a; Soch et al., in prep.; Richter et al., submitted). One potential explanation for this finding may be that, in our study, the sample investigated consisted of neurologically and psychiatrically healthy older adults without signs of cognitive impairment. This suggests that brain atrophy (i.e., structural volume loss) may to some extent occur invariably with increasing age, but does not necessarily affect cognitive performance as long as (i) the degree is still within the bounds of normal aging and (ii) it is not accompanied by functional processing changes (reflected in fMRI scores), potentially due to compensatory mechanisms (Kizilirmak et al., 2021). This is in line with previous studies that reported a decoupling between gray and white matter measures and memory performance in older age (de Mooij et al., 2018), underscoring that cognitive maintenance or reserve is – at least to a degree – independent of neural maintenance. A large meta-analysis also highlights the lack of a strong dependency between structural and cognitive decline (Oschwald et al., 2019), suggesting that the healthy aging brain possesses a considerable potential to compensate for inevitable age-related structural decline (Stern, 2009; Nyberg et al., 2012; Cabeza et al., 2018).

### 4.5. Conclusion

We have shown a systematic difference in predictive ability between structural MRI markers (and resting-state fMRI) on the one hand versus functional MRI markers (especially fMRI summary statistics) on the other hand. Whereas the former are most strongly related to chronological age reflecting the mere progression of time, the latter allow to better predict cognitive performance in episodic memory. In a sense, this double dissociation supports the concept of cognitive reserve as a phenomenon that may to some degree be independent from structural brain aging. Further research has to elucidate the sources of preserved memory performance in older adults with structural degradation, but functional maintenance.

## Footnotes

1 URL: https://github.com/JoramSoch/ML4ML.

2 Confidence intervals were generated using the MATLAB functions *binofit* for accuracies (assuming that the numbers of correct predictions are binomially distributed with unknown success probability) and *corrcoef* for correlations (assuming that actual and predicted continuous variables are linearly related).

3 Note that we are here not using behavioral data as source variables, because the target variable of memory performance is a mathematical function of the behavioral response frequencies. For this reason, prediction from response frequencies to all items would reach ceiling performance and is not shown.

## References

Ackenheil, M., Stotz, G., Dietz-Bauer, R., Vossen, A., 1999. Mini International Neuropsychiatric Interview - German version 5.0.0.

Assmann, A., Richter, A., Schütze, H., Soch, J., Barman, A., Behnisch, G., Knopf, L., Raschick, M., Schult, A., Wüstenberg, T., Behr, J., Düzel, E., Seidenbecher, C.I., Schott, B.H., 2020. Neurocan genome-wide psychiatric risk variant affects explicit memory performance and hippocampal function in healthy humans. Eur J Neurosci ejn.14872. https://doi.org/10.1111/ejn.14872

Bagarinao, E., Watanabe, H., Maesawa, S., Kawabata, K., Hara, K., Ohdake, R., Ogura, A., Mori, D., Yoneyama, N., Imai, K., Yokoi, T., Kato, T., Koyama, S., Katsuno, M., Wakabayashi, T., Kuzuya, M., Hoshiyama, M., Isoda, H., Naganawa, S., Ozaki, N., Sobue, G., 2022. Reserve and Maintenance in the Aging Brain: A Longitudinal Study of Healthy Older Adults. eNeuro 9, ENEURO.0455-21.2022. https://doi.org/10.1523/ENEURO.0455-21.2022

Bainbridge, W.A., Berron, D., Schütze, H., Cardenas-Blanco, A., Metzger, C., Dobisch, L., Bittner, D., Glanz, W., Spottke, A., Rudolph, J., Brosseron, F., Buerger, K., Janowitz, D., Fliessbach, K., Heneka, M., Laske, C., Buchmann, M., Peters, O., Diesing, D., Li, S., Priller, J., Spruth, E.J., Altenstein, S., Schneider, A., Kofler, B., Teipel, S., Kilimann, I., Wiltfang, J., Bartels, C., Wolfsgruber, S., Wagner, M., Jessen, F., Baker, C.I., Düzel, E., 2019. Memorability of photographs in subjective cognitive decline and mild cognitive impairment: Implications for cognitive assessment. Alzheimer’s & Dementia: Diagnosis, Assessment & Disease Monitoring 11, 610–618. https://doi.org/10.1016/j.dadm.2019.07.005

Barman, A., Assmann, A., Richter, S., Soch, J., Schütze, H., Wüstenberg, T., Deibele, A., Klein, M., Richter, A., Behnisch, G., Düzel, E., Zenker, M., Seidenbecher, C.I., Schott, B.H., 2014. Genetic variation of the RASGRF1 regulatory region affects human hippocampus-dependent memory. Front. Hum. Neurosci. 8. https://doi.org/10.3389/fnhum.2014.00260

Bashyam, V.M., Erus, G., Doshi, J., Habes, M., Nasrallah, I.M., Truelove-Hill, M., Srinivasan, D., Mamourian, L., Pomponio, R., Fan, Y., Launer, L.J., Masters, C.L., Maruff, P., Zhuo, C., Völzke, H., Johnson, S.C., Fripp, J., Koutsouleris, N., Satterthwaite, T.D., Wolf, D., Gur, R.E., Gur, R.C., Morris, J., Albert, M.S., Grabe, H.J., Resnick, S., Bryan, R.N., Wolk, D.A., Shou, H., Davatzikos, C., 2020. MRI signatures of brain age and disease over the lifespan based on a deep brain network and 14 468 individuals worldwide. Brain 143, 2312–2324. https://doi.org/10.1093/brain/awaa160

Billette, O., Ziegler, G., Düzel, E., Maass, A., 2022. Novelty-related fMRI responses of precuneus and medial temporal regions in individuals at risk for Alzheimer’s disease.

Brodersen, K.H., Ong, C.S., Stephan, K.E., Buhmann, J.M., 2010. The Balanced Accuracy and Its Posterior Distribution, in: 2010 20th International Conference on Pattern Recognition. Presented at the 2010 20th International Conference on Pattern Recognition (ICPR), IEEE, Istanbul, Turkey, pp. 3121–3124. https://doi.org/10.1109/ICPR.2010.764

Buckner, R.L., Andrews-Hanna, J.R., Schacter, D.L., 2008. The Brain’s Default Network: Anatomy, Function, and Relevance to Disease. Annals of the New York Academy of Sciences 1124, 1–38. https://doi.org/10.1196/annals.1440.011

Cabeza, R., Albert, M., Belleville, S., Craik, F.I.M., Duarte, A., Grady, C.L., Lindenberger, U., Nyberg, L., Park, D.C., Reuter-Lorenz, P.A., Rugg, M.D., Steffener, J., Rajah, M.N., 2018. Maintenance, reserve and compensation: the cognitive neuroscience of healthy ageing. Nature Reviews Neuroscience 19, 701–710. https://doi.org/10.1038/s41583-018-0068-2

Cabeza, R., Nyberg, L., Park, D., 2004. Cognitive Neuroscience of Aging. Oxford University Press. https://doi.org/10.1093/acprof:oso/9780195156744.001.0001

Cole, J.H., Franke, K., 2017. Predicting Age Using Neuroimaging: Innovative Brain Ageing Biomarkers. Trends in Neurosciences 40, 681–690. https://doi.org/10.1016/j.tins.2017.10.001

Cole, J.H., Poudel, R.P.K., Tsagkrasoulis, D., Caan, M.W.A., Steves, C., Spector, T.D., Montana, G., 2017. Predicting brain age with deep learning from raw imaging data results in a reliable and heritable biomarker. NeuroImage 163, 115–124. https://doi.org/10.1016/j.neuroimage.2017.07.059

Cole, J.H., Ritchie, S.J., Bastin, M.E., Valdés Hernández, M.C., Muñoz Maniega, S., Royle, N., Corley, J., Pattie, A., Harris, S.E., Zhang, Q., Wray, N.R., Redmond, P., Marioni, R.E., Starr, J.M., Cox, S.R., Wardlaw, J.M., Sharp, D.J., Deary, I.J., 2018. Brain age predicts mortality. Mol Psychiatry 23, 1385–1392. https://doi.org/10.1038/mp.2017.62

de Mooij, S.M.M., Henson, R.N.A., Waldorp, L.J., Kievit, R.A., 2018. Age Differentiation within Gray Matter, White Matter, and between Memory and White Matter in an Adult Life Span Cohort. The Journal of Neuroscience 38, 5826–5836. https://doi.org/10.1523/JNEUROSCI.1627-17.2018

Dounavi, M.-E., Mak, E., Wells, K., Ritchie, K., Ritchie, C.W., Su, L., O’ Brien, J.T., 2020. Volumetric alterations in the hippocampal subfields of subjects at increased risk of dementia. Neurobiology of Aging 91, 36–44. https://doi.org/10.1016/j.neurobiolaging.2020.03.006

Duarte, A., Graham, K.S., Henson, R.N., 2010. Age-related changes in neural activity associated with familiarity, recollection and false recognition. Neurobiology of Aging 31, 1814–1830. https://doi.org/10.1016/j.neurobiolaging.2008.09.014

Düzel, E., Acosta-Cabronero, J., Berron, D., Biessels, G.J., Björkman-Burtscher, I., Bottlaender, M., Bowtell, R., Buchem, M. v., Cardenas-Blanco, A., Boumezbeur, F., Chan, D., Clare, S., Costagli, M., Rochefort, L., Fillmer, A., Gowland, P., Hansson, O., Hendrikse, J., Kraff, O., Ladd, M.E., Ronen, I., Petersen, E., Rowe, J.B., Siebner, H., Stoecker, T., Straub, S., Tosetti, M., Uludag, K., Vignaud, A., Zwanenburg, J., Speck, O., 2019. European Ultrahigh-Field Imaging Network for Neurodegenerative Diseases (EUFIND). Alzheimer’s & Dementia: Diagnosis, Assessment & Disease Monitoring 11, 538–549. https://doi.org/10.1016/j.dadm.2019.04.010

Düzel, E., Berron, D., Schütze, H., Cardenas-Blanco, A., Metzger, C., Betts, M., Ziegler, G., Chen, Y., Dobisch, L., Bittner, D., Glanz, W., Reuter, M., Spottke, A., Rudolph, J., Brosseron, F., Buerger, K., Janowitz, D., Fliessbach, K., Heneka, M., Laske, C., Buchmann, M., Nestor, P., Peters, O., Diesing, D., Li, S., Priller, J., Spruth, E.J., Altenstein, S., Ramirez, A., Schneider, A., Kofler, B., Speck, O., Teipel, S., Kilimann, I., Dyrba, M., Wiltfang, J., Bartels, C., Wolfsgruber, S., Wagner, M., Jessen, F., 2018. CSF total tau levels are associated with hippocampal novelty irrespective of hippocampal volume. Alzheimer’s & Dementia: Diagnosis, Assessment & Disease Monitoring 10, 782–790. https://doi.org/10.1016/j.dadm.2018.10.003

Düzel, E., Schütze, H., Yonelinas, A.P., Heinze, H.-J., 2010. Functional phenotyping of successful aging in long-term memory: Preserved performance in the absence of neural compensation. Hippocampus 803–814. https://doi.org/10.1002/hipo.20834

Dyrba, M., Hanzig, M., Altenstein, S., Bader, S., Ballarini, T., Brosseron, F., Buerger, K., Cantré, D., Dechent, P., Dobisch, L., Düzel, E., Ewers, M., Fliessbach, K., Glanz, W., Haynes, J.-D., Heneka, M.T., Janowitz, D., Keles, D.B., Kilimann, I., Laske, C., Maier, F., Metzger, C.D., Munk, M.H., Perneczky, R., Peters, O., Preis, L., Priller, J., Rauchmann, B., Roy, N., Scheffler, K., Schneider, A., Schott, B.H., Spottke, A., Spruth, E.J., Weber, M.-A., Ertl-Wagner, B., Wagner, M., Wiltfang, J., Jessen, F., Teipel, S.J., for the ADNI, AIBL, DELCODE study groups, 2021. Improving 3D convolutional neural network comprehensibility via interactive visualization of relevance maps: evaluation in Alzheimer’s disease. Alz Res Therapy 13, 191. https://doi.org/10.1186/s13195-021-00924-2

Eitel, F., Schulz, M.-A., Seiler, M., Walter, H., Ritter, K., 2021. Promises and pitfalls of deep neural networks in neuroimaging-based psychiatric research. Experimental Neurology 339, 113608. https://doi.org/10.1016/j.expneurol.2021.113608

Elliott, M.L., Knodt, A.R., Ireland, D., Morris, M.L., Poulton, R., Ramrakha, S., Sison, M.L., Moffitt, T.E., Caspi, A., Hariri, A.R., 2020. What Is the Test-Retest Reliability of Common Task-Functional MRI Measures? New Empirical Evidence and a Meta-Analysis. Psychol Sci 31, 792–806. https://doi.org/10.1177/0956797620916786

Fisch, L., Leenings, R., Winter, N.R., Dannlowski, U., Gaser, C., Cole, J.H., Hahn, T., 2021. Editorial: Predicting Chronological Age From Structural Neuroimaging: The Predictive Analytics Competition 2019. Front. Psychiatry 12, 710932. https://doi.org/10.3389/fpsyt.2021.710932

Fjell, A.M., McEvoy, L., Holland, D., Dale, A.M., Walhovd, K.B., for the Alzheimer’s Disease Neuroimaging Initiative, 2013. Brain Changes in Older Adults at Very Low Risk for Alzheimer’s Disease. Journal of Neuroscience 33, 8237–8242. https://doi.org/10.1523/JNEUROSCI.5506-12.2013

Fjell, A.M., Walhovd, K.B., Fennema-Notestine, C., McEvoy, L.K., Hagler, D.J., Holland, D., Brewer, J.B., Dale, A.M., 2009. One-Year Brain Atrophy Evident in Healthy Aging. Journal of Neuroscience 29, 15223–15231. https://doi.org/10.1523/JNEUROSCI.3252-09.2009

Foo, H., Thalamuthu, A., Jiang, J., Koch, F., Mather, K.A., Wen, W., Sachdev, P.S., 2021. Age-and Sex-Related Topological Organization of Human Brain Functional Networks and Their Relationship to Cognition. Front. Aging Neurosci. 13, 758817. https://doi.org/10.3389/fnagi.2021.758817

Gong, W., Beckmann, C.F., Vedaldi, A., Smith, S.M., Peng, H., 2021. Optimising a Simple Fully Convolutional Network for Accurate Brain Age Prediction in the PAC 2019 Challenge. Front. Psychiatry 12, 627996. https://doi.org/10.3389/fpsyt.2021.627996

Habes, M., Pomponio, R., Shou, H., Doshi, J., Mamourian, E., Erus, G., Nasrallah, I., Launer, L.J., Rashid, T., Bilgel, M., Fan, Y., Toledo, J.B., Yaffe, K., Sotiras, A., Srinivasan, D., Espeland, M., Masters, C., Maruff, P., Fripp, J., Völzk, H., Johnson, S.C., Morris, J.C., Albert, M.S., Miller, M.I., Bryan, R.N., Grabe, H.J., Resnick, S.M., Wolk, D.A., Davatzikos, C., for the iSTAGING consortium, the Preclinical AD consortium, the ADNI, and the CARDIA studies, 2021. The Brain Chart of Aging: Machine-learning analytics reveals links between brain aging, white matter disease, amyloid burden, and cognition in the iSTAGING consortium of 10,216 harmonized MR scans. Alzheimer’s & Dementia 17, 89–102. https://doi.org/10.1002/alz.12178

Härting, C., Markowitsch, H.-J., Neufeld, H., Calabrese, P., Deisinger, K., Kessler, J., 2000. Wechsler Memory Scale, Revised Edition, German Edition. ed. Huber, Bern.

Helmstaedter, C., Lendt, M., Lux, S., 2001. Verbaler Lern-und Merkfähigkeitstest, 1st ed. Beltz Test GmbH, Göttingen, Germany.

Hinrichs, H., Scholz, M., Tempelmann, C., Woldorff, M.G., Dale, A.M., Heinze, H.-J., 2000. Deconvolution of Event-Related fMRI Responses in Fast-Rate Experimental Designs: Tracking Amplitude Variations. Journal of Cognitive Neuroscience 12, 76–89. https://doi.org/10.1162/089892900564082

Jessen, F., Spottke, A., Boecker, H., Brosseron, F., Buerger, K., Catak, C., Fliessbach, K., Franke, C., Fuentes, M., Heneka, M.T., Janowitz, D., Kilimann, I., Laske, C., Menne, F., Nestor, P., Peters, O., Priller, J., Pross, V., Ramirez, A., Schneider, A., Speck, O., Spruth, E.J., Teipel, S., Vukovich, R., Westerteicher, C., Wiltfang, J., Wolfsgruber, S., Wagner, M., Düzel, E., 2018. Design and first baseline data of the DZNE multicenter observational study on predementia Alzheimer’s disease (DELCODE). Alzheimer’s Research and Therapy 10, 1–10. https://doi.org/10.1186/s13195-017-0314-2

Jia, X.-Z., Sun, J.-W., Ji, G.-J., Liao, W., Lv, Y.-T., Wang, J., Wang, Z., Zhang, H., Liu, D.-Q., Zang, Y.-F., 2020. Percent amplitude of fluctuation: A simple measure for resting-state fMRI signal at single voxel level. PLoS ONE 15, e0227021.https://doi.org/10.1371/journal.pone.0227021

Jia, X.-Z., Wang, J., Sun, H.-Y., Zhang, H., Liao, W., Wang, Z., Yan, C.-G., Song, X.-W., Zang, Y.-F., 2019. RESTplus: an improved toolkit for resting-state functional magnetic resonance imaging data processing. Science Bulletin 64, 953–954. https://doi.org/10.1016/j.scib.2019.05.008

Kizilirmak, J.M., 2021. Age-dependent involvement of default mode network structures in episodic long-term memory formation. OSF Retrieved from osf.io/gfw85.

Kizilirmak, J.M., Fischer, L., Krause, J., Soch, J., Richter, A., Schott, B.H., 2021. Learning by Insight-Like Sudden Comprehension as a Potential Strategy to Improve Memory Encoding in Older Adults. Front. Aging Neurosci. 13, 661346. https://doi.org/10.3389/fnagi.2021.661346

Luders, E., Cherbuin, N., Gaser, C., 2016. Estimating brain age using high-resolution pattern recognition: Younger brains in long-term meditation practitioners. NeuroImage 134, 508–513. https://doi.org/10.1016/j.neuroimage.2016.04.007

Maillet, D., Rajah, M.N., 2014. Age-related differences in brain activity in the subsequent memory paradigm: A meta-analysis. Neuroscience & Biobehavioral Reviews 45, 246–257. https://doi.org/10.1016/j.neubiorev.2014.06.006

Mevel, K., Chételat, G., Eustache, F., Desgranges, B., 2011. The Default Mode Network in Healthy Aging and Alzheimer’s Disease. International Journal of Alzheimer’s Disease 2011, 1–9. https://doi.org/10.4061/2011/535816

Minkova, L., Habich, A., Peter, J., Kaller, C.P., Eickhoff, S.B., Klöppel, S., 2017. Gray matter asymmetries in aging and neurodegeneration: A review and meta-analysis: VBM-ALE Analysis of GM Asymmetries. Hum. Brain Mapp. 38, 5890–5904. https://doi.org/10.1002/hbm.23772

Naismith, S.L., Glozier, N., Burke, D., Carter, P.E., Scott, E., Hickie, I.B., 2009. Early intervention for cognitive decline: Is there a role for multiple medical or behavioural interventions? Early Intervention in Psychiatry 3, 19–27. https://doi.org/10.1111/j.1751-7893.2008.00102.x

Nyberg, L., Lövdén, M., Riklund, K., Lindenberger, U., Bäckman, L., 2012. Memory aging and brain maintenance. Trends in Cognitive Sciences 16, 292–305. https://doi.org/10.1016/j.tics.2012.04.005

Nyberg, L., Pudas, S., 2019. Successful Memory Aging. Annu. Rev. Psychol. 70, 219–243. https://doi.org/10.1146/annurev-psych-010418-103052

Oschwald, J., Guye, S., Liem, F., Rast, P., Willis, S., Röcke, C., Jäncke, L., Martin, M., Mérillat, S., 2019. Brain structure and cognitive ability in healthy aging: a review on longitudinal correlated change. Reviews in the Neurosciences 31, 1–57. https://doi.org/10.1515/revneuro-2018-0096

Richter, A., Soch, J., Kizilirmak, J.M., Fischer, L., Schütze, H., Assmann, A., Behnisch, G., Feldhoff, H., Knopf, L., Raschick, M., Schult, A., Seidenbecher, C.I., Yakupov, R., Düzel, E., Schott, B.H., 2022. Summary statistics of memory-related fMRI activity reflect dissociable neuropsychological and anatomical signatures of neurocognitive aging (preprint). bioRxiv Neuroscience. https://doi.org/10.1101/2022.02.04.479169

Scarmeas, N., Zarahn, E., Anderson, K.E., Hilton, J., Flynn, J., Van Heertum, R.L., Sackeim, H.A., Stern, Y., 2003. Cognitive reserve modulates functional brain responses during memory tasks: a PET study in healthy young and elderly subjects. NeuroImage 19, 1215–1227. https://doi.org/10.1016/S1053-8119(03)00074-0

Schiltz, K., Szentkuti, A., Guderian, S., Kaufmann, J., Münte, T.F., Heinze, H.J., Düzel, E., 2006. Relationship between Hippocampal Structure and Memory Function in Elderly Humans. Journal of Cognitive Neuroscience 18, 990–1003. https://doi.org/10.1162/jocn.2006.18.6.990

Schott, B.H., Assmann, A., Schmierer, P., Soch, J., Erk, S., Garbusow, M., Mohnke, S., Pöhland, L., Romanczuk-Seiferth, N., Barman, A., Wüstenberg, T., Haddad, L., Grimm, O., Witt, S., Richter, S., Klein, M., Schütze, H., Mühleisen, T.W., Cichon, S., Rietschel, M., Noethen, M.M., Tost, H., Gundelfinger, E.D., Düzel, E., Heinz, A., Meyer-Lindenberg, A., Seidenbecher, C.I., Walter, H., 2014. Epistatic interaction of genetic depression risk variants in the human subgenual cingulate cortex during memory encoding. Transl Psychiatry 4, e372–e372. https://doi.org/10.1038/tp.2014.10

Sheehan, D.V., Lecrubier, Y., Sheehan, H., Amorim, P., Janavs, J., Weiller, E., Hergueta, T., Baker, R., Dunbar, G.C., 1998. The Mini-International Neuropsychiatric Interview (M.I.N.I.): the development and validation of a structured diagnostic psychiatric interview for DSM-IV and ICD-10. Journal of Clinical Psychiatry 59, 22–33.

Soch, J., 2020. Distributional Transformation Improves Decoding Accuracy When Predicting Chronological Age from Structural MRI. Front. Psychiatry 11, 604268. https://doi.org/10.3389/fpsyt.2020.604268

Soch, J., Richter, A., Schott, B.H., Kizilirmak, J.M., 2022. A novel approach for modelling combined recognition-confidence ratings by separating decidedness, recognition and confidence.

Soch, J., Richter, A., Schütze, H., Kizilirmak, J.M., Assmann, A., Behnisch, G., Feldhoff, H., Fischer, L., Heil, J., Knopf, L., Merkel, C., Raschick, M., Schietke, C., Schult, A., Seidenbecher, C.I., Yakupov, R., Ziegler, G., Wiltfang, J., Düzel, E., Schott, B.H., 2021a. A comprehensive score reflecting memory-related fMRI activations and deactivations as potential biomarker for neurocognitive aging. Hum Brain Mapp 42, 4478–4496. https://doi.org/10.1002/hbm.25559

Soch, J., Richter, A., Schütze, H., Kizilirmak, J.M., Assmann, A., Knopf, L., Raschick, M., Schult, A., Maass, A., Ziegler, G., Richardson-Klavehn, A., Düzel, E., Schott, B.H., 2021b. Bayesian model selection favors parametric over categorical fMRI subsequent memory models in young and older adults. NeuroImage 230, 117820. https://doi.org/10.1016/j.neuroimage.2021.117820

Steffener, J., Habeck, C., O’Shea, D., Razlighi, Q., Bherer, L., Stern, Y., 2016. Differences between chronological and brain age are related to education and self-reported physical activity. Neurobiology of Aging 40, 138–144. https://doi.org/10.1016/j.neurobiolaging.2016.01.014

Stern, Y., 2012. Cognitive reserve in ageing and Alzheimer’s disease. The Lancet Neurology 11, 1006–1012. https://doi.org/10.1016/S1474-4422(12)70191-6

Stern, Y., 2009. Cognitive reserve. Neuropsychologia 47, 2015–2028. https://doi.org/10.1016/j.neuropsychologia.2009.03.004

Veldsman, M., Nobis, L., Alfaro-Almagro, F., Manohar, S., Husain, M., 2021. The human hippocampus and its subfield volumes across age, sex and APOE e4 status. Brain Communications 3, fcaa219. https://doi.org/10.1093/braincomms/fcaa219

Whitty, E., Mansour, H., Aguirre, E., Palomo, M., Charlesworth, G., Ramjee, S., Poppe, M., Brodaty, H., Kales, H.C., Morgan-Trimmer, S., Nyman, S.R., Lang, I., Walters, K., Petersen, I., Wenborn, J., Minihane, A.-M., Ritchie, K., Huntley, J., Walker, Z., Cooper, C., 2020. Efficacy of lifestyle and psychosocial interventions in reducing cognitive decline in older people: Systematic review. Ageing Research Reviews 62, 101113. https://doi.org/10.1016/j.arr.2020.101113

Woodard, J.L., Seidenberg, M., Nielson, K.A., Smith, J.C., Antuono, P., Durgerian, S., Guidotti, L., Zhang, Q., Butts, A., Hantke, N., Lancaster, M., Rao, S.M., 2010. Prediction of cognitive decline in healthy older adults using fMRI. Journal of Alzheimer’s Disease 21, 871–885. https://doi.org/10.3233/JAD-2010-091693

Xing, X.-X., 2021. Globally Aging Cortical Spontaneous Activity Revealed by Multiple Metrics and Frequency Bands Using Resting-State Functional MRI. Front. Aging Neurosci. 13, 803436. https://doi.org/10.3389/fnagi.2021.803436

