## Supplementary Material for "Structural and functional MRI data differentially predict chronological age and behavioral memory performance"

[1] German Center for Neurodegenerative Diseases (DZNE), Göttingen, Germany
[2] Bernstein Center for Computational Neuroscience (BCCN), Berlin, Germany
[3] Leibniz Institute for Neurobiology (LIN), Magdeburg, Germany
[4] German Center for Neurodegenerative Diseases (DZNE), Magdeburg, Germany
[5] Otto von Guericke University, Medical Faculty, Magdeburg, Germany
[6] Center for Behavioral Brain Sciences (CBBS), Magdeburg, Germany
[7] Department of Psychiatry and Psychotherapy, University Medical Center Göttingen, Göttingen, Germany
* These authors contributed equally to this work.

### Supplementary Methods

##### Partitioning of older subjects for post-hoc analyses

In order to perform group comparisons following the predictive analyses for age and memory (see Sections 3.1 and 3.2 in the main paper), older subjects were split into four groups based on the quartiles of chronological age and memory performance, respectively. Data from subjects outside the interquartile range, i.e. below the first quartile or above the third quartile (see Figure S3), were used for post-hoc analyses of functional and structural MRI (see Figure 5).

### Supplementary Results

##### Prediction of independent memory performance in young subjects

Independent measures of memory performance (see Section 2.6 and Table 2 in the main paper) could not be reliably predicted from measured data in young subjects (see Figure S2). This is in contrast to the older subjects for whom we observed robust above-chance predictive correlations for some feature sets (see Figure 4).

### Supplementary Discussion

##### Differences in predictive accuracy for independent memory performance

When predicting outcomes in VLMT and WMS memory tests (see Section 3.3 and Figure 4 in the main paper), we observed that VLMT performance could be predicted from behavioral responses to old items of the fMRI memory task (FADE), but not from task-based fMRI contrast maps, and the reverse was observed for WMS performance.

This is presumably because the verbal-semantic VLMT includes a distractor list and the distractors act similar like the new items in the FADE task, requiring subjects to decide during item retrieval, whether an item they remember was in the target list or the distractor list. This similarity of discrimination requirements might induce a correlation between the number of old items recalled (VLMT) and the fraction of old images recognized (FADE), leading to a significant predictive correlation. This interpretation would be in line a two-process model for recognition and retrieval (Anderson & Bower, 1972) which points out the importance of contextual information, e.g. distractor lists during learning (Cox & Dobbins, 2011).

### Supplementary Figures

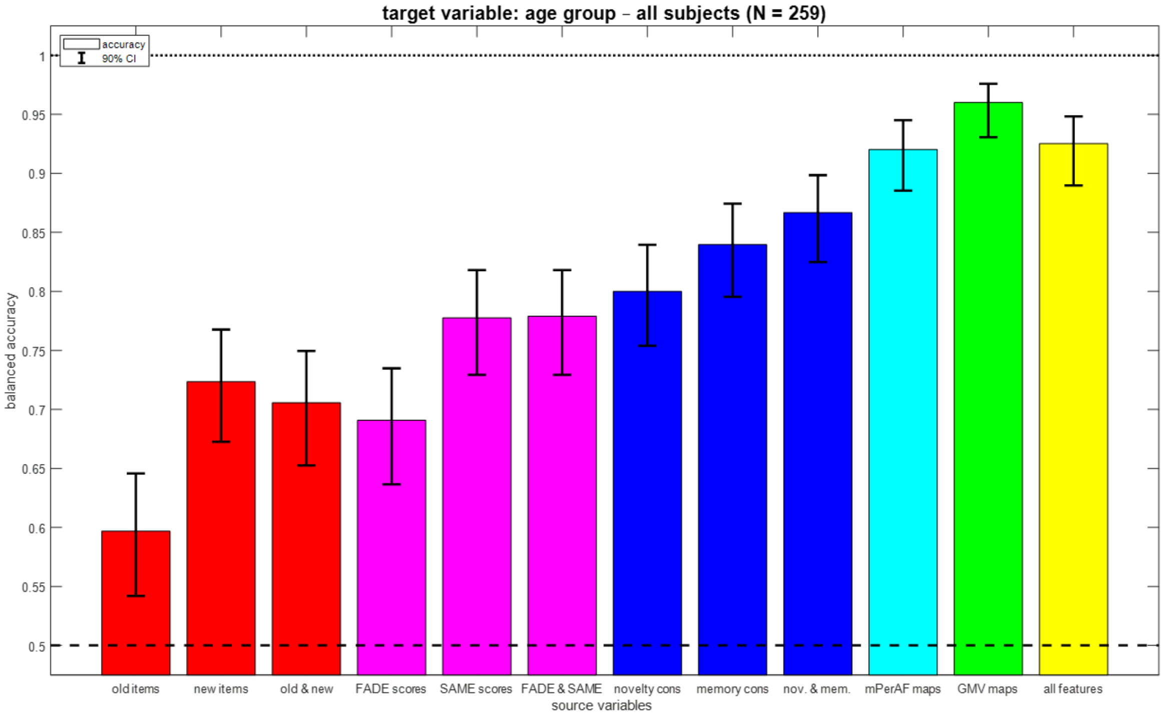


**Figure S1.** *Classification of age group from different feature sets.* Bar plots show accuracy for decoding age group (young vs. older) from behavioral data (red), fMRI scores (magenta), task-based fMRI contrasts (blue), resting-state fMRI maps and structural MRI (green), or all features (yellow). Error bars denote 90% confidence intervals; chance level and ceiling performance are indicated by dashed and dotted lines, respectively; x-axis labels are explained in Table 4.

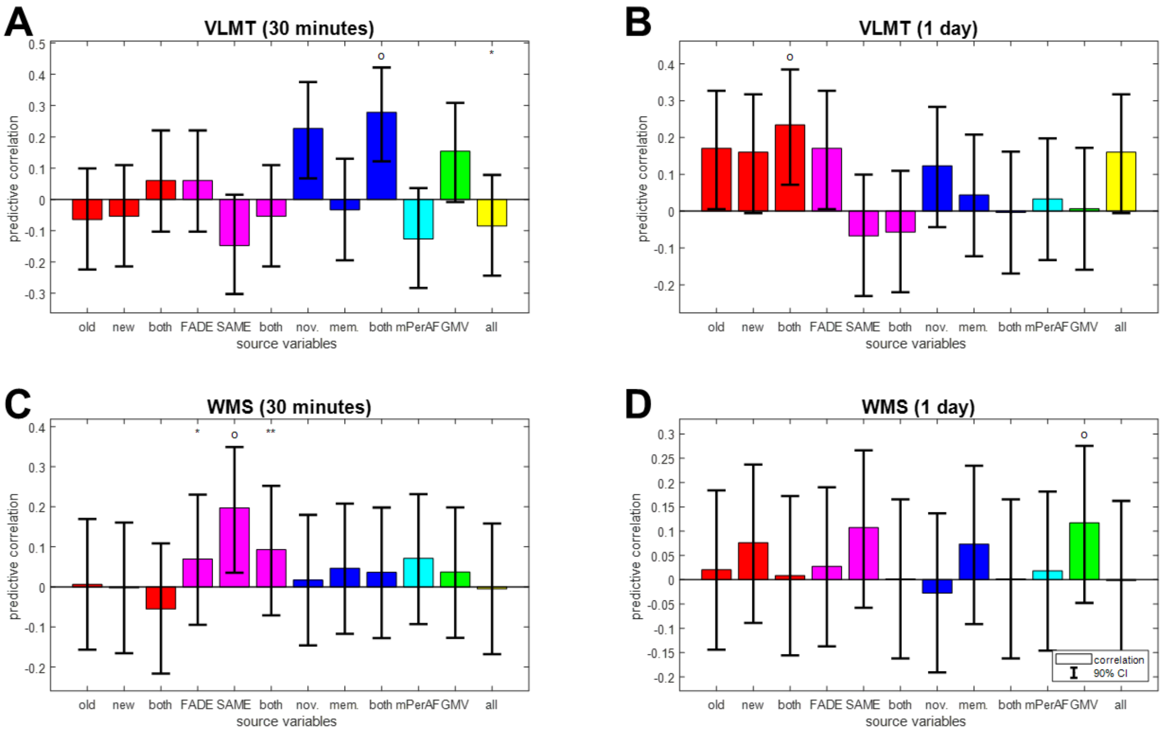


**Figure S2.** *Reconstruction of independent memory performance in young subjects.* Bar plots show correlation coefficients for predicting, in young subjects only, independent measures of memory performance, namely **(A)** VLMT items after 30 minutes, **(B)** VLMT items after 1 day, **(C)** WMS points after 30 minutes and **(D)** WMS points after 1 day, from behavioral data (red), fMRI scores (magenta), task-based fMRI contrasts (blue), resting-state fMRI maps (cyan) and structural MRI (green), or all features (yellow). This figure mirrors Figure 4 from the main paper. The layout follows that of Figure 3C from the main paper.

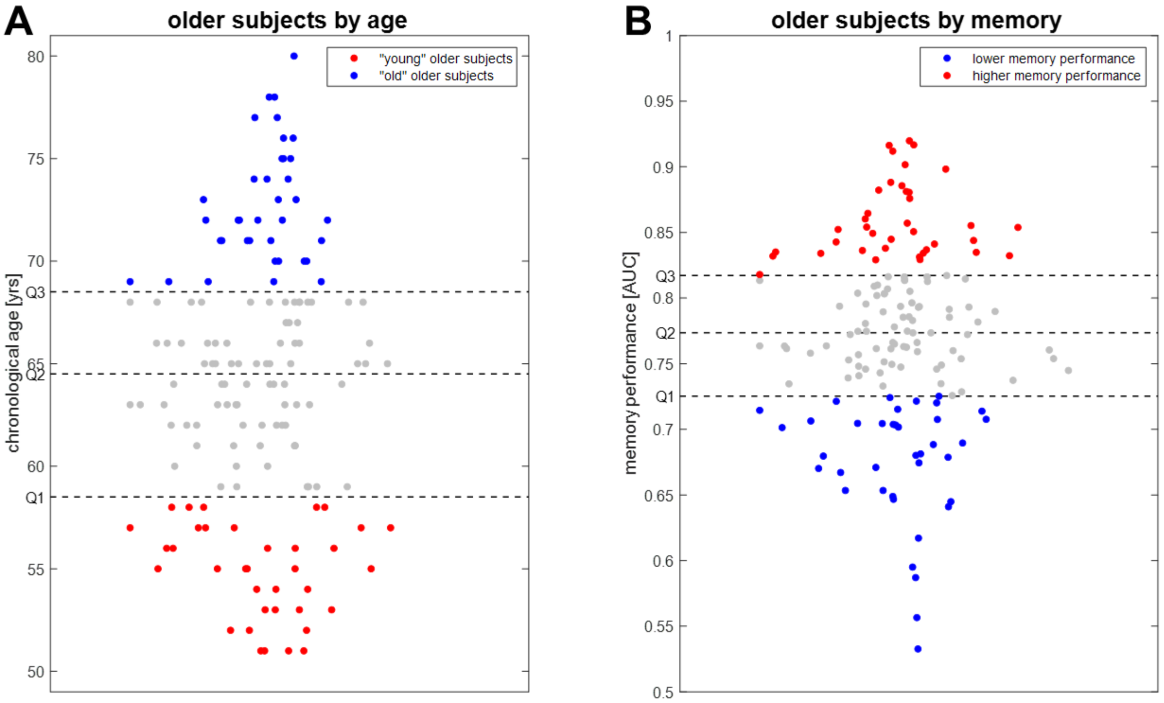


**Figure S3.** *Separation of older subjects by chronological age and memory performance.* Older subjects were partitioned into four groups based on quartiles (dashed black lines) obtained from the observed distributions of **(A)** chronological age and **(B)** memory performance. Subjects at the lower and the upper end (red and blue) were used for post-hoc analyses reported in Sections 3.4 and 3.5 of the main paper.

#### References

Anderson, J.R., Bower, G.H., 1972. Recognition and retrieval processes in free recall. Psychological Review 79, 97–123. https://doi.org/10.1037/h0033773

Cox, J.C., Dobbins, I.G., 2011. The striking similarities between standard, distractor-free, and target-free recognition. Memory & Cognition 39, 925–940. https://doi.org/10.3758/s13421-011-0090-3
